# Ultrastructural analysis of SARS-CoV-2 interactions with the host cell via high resolution scanning electron microscopy

**DOI:** 10.1101/2020.06.10.144642

**Authors:** Lucio Ayres Caldas, Fabiana Avila Carneiro, Luiza Mendonça Higa, Fábio Luiz Monteiro, Gustavo Peixoto da Silva, Luciana Jesus da Costa, Edison Luiz Durigon, Amilcar Tanuri, Wanderley de Souza

**Author notes:** **Corresponding author:** Laboratório de Ultraestrutura Celular Hertha Meyer. Av. Carlos Chagas Filho 373, Prédio CCS, Bloco C, subsolo, CEP:21941902, Cidade Universitária, Rio de Janeiro, RJ, Brazil. Present/Permanent address: Núcleo Multidisciplinar de Pesquisa UFRJ-Xerém em Biologia – NUMPEX-BIO, Universidade Federal do Rio de Janeiro, Campus Duque de Caxias Geraldo Cidade. CEP: 25265-970, Rio de Janeiro, RJ, Brazil.

## Abstract

SARS-CoV-2 is the cause of the ongoing COVID-19 pandemic. Here, we investigated the interaction of this new coronavirus with Vero cells using high resolution scanning electron microscopy. Surface morphology, the interior of infected cells and the distribution of viral particles in both environments were observed 2 and 48 hours after infection. We showed areas of viral processing, details of vacuole contents, and viral interactions with the cell surface. Intercellular connections were also approached, and viral particles were adhered to these extensions suggesting direct cell-to-cell transmission of SARS-CoV-2.

**Highlights:** We used high resolution scanning electron microscopy to investigate Vero cells infected with SARS-CoV-2 at 2 and 48 hours post-infection. The central conclusions of this work include:

- Infected cells display polarization of their cytosol forming a restricted viroplasm-like zone dedicated to virus production and morphogenesis.
- This is the first demonstration of SARS-CoV-2 attachment by scanning electron microscopy.
- This is the first scanning electron microscopy images of the interior of SARS-CoV-2 infected cells and exploration of their vacuole contents.
- Perspective-viewing of bordering vesicles in close association with vacuoles.
- Observation of membrane ruffles and structures suggestive of exocytosis on the surface of infected cells.
- The first demonstration of viral surfing in cell-to-cell communication on SARS-CoV-2 infection.

## 1. Introduction

COVID-19 is an acute respiratory illness caused by the SARS-CoV-2—a novel coronavirus identified during this pandemic [1]. The outbreak started at Wuhan in Hubei province, China, in December 2019 [2]. Since then, the world has seen a rapid spread of the virus with an increasing number of infected people—around 6 million cases and close to 400,000 deaths [3]. In the first four months, the outbreak has led to more than 28,000 deaths in Brazil [4]. There is currently no vaccine or specific treatment for COVID-19. Patients attendance is mainly based on supportive and symptomatic care. Therefore, a treatment capable of inhibiting viral infection and/or replication is urgent.

SARS-CoV-2 is an enveloped, positive-sense RNA beta-Coronavirus belonging to the *Coronaviridae* family. The genome is packaged inside a helix capsid formed by the nucleocapsid protein (N). Three other structural proteins are associated with the viral envelope: membrane (M), envelope (E), and glycoprotein spike (S). Cellular entry of SARS-CoV-2 depends on the binding of the S protein to angiotensin converting enzyme 2 (ACE2)—a specific cellular receptor located on the surface of the host cell [5,6]. This is a common receptor for SARS-CoV as well [7,8] (Li et al., 2003; 2005); this receptor facilitates zoonotic transfer because these viruses can engage ACE2 from different animal species [9].

Beta-coronaviruses replicate in the cytoplasm; cellular compartments like the endoplasmic reticulum (ER) and the endoplasmic reticulum-Golgi apparatus intermediate compartiment (ERGIC) go through intense remodeling. This implies the contribution of host membranes and organelles for viral replication. Therefore, remodeling of intracellular membranes due to coronavirus infection is also observed for many RNA viruses [10].

After internalization and RNA release into the cytoplasm, a set of proteins is synthesized triggering the formation of vesicles that become a viral platform ensuring efficient replication and transcription of the RNA [11,12].

New coronavirus particles are assembled in the endoplasmic reticulum and Golgi complex. Membrane budding between these compartments was reported in association with N protein and genomic RNA along with M, E, and S proteins. The complete virions are delivered to the extracellular environment following a conventional secretory route [13–15].

The research community has sought to better understand the genetic makeup of the virus and thus discover how to effectively treat it. Social isolation for 14 days is the main way to prevent the disease from spreading. Quarantine and lockdowns were implemented in cities with high rates of infection and mortality [3]. Death is common in patients with severe symptoms including shortness of breathing, fever, lethargy, respiratory failure, and/or thrombosis [16,17].

Understanding the virus-cell interactions is key to vaccines, treatments, and diagnoses. Most microscopic studies of SARS-CoV-2 were performed with transmission electron microscopy. Here, we used high resolution scanning electron microscopy (SEM) to study inner cellular structures. The results offer evidence of infection-induced cellular remodeling and the formation of a specialized region for viral morphogenesis. We also show intercellular extensions for viral cell surfing. These observations offer new insights into the transmission of SARS-CoV-2.

## 2. Material and Methods

### Cells and Virus

SARS-CoV-2 isolate (HIAE-02: SARS-CoV-2/SP02/human/2020/BRA (GenBank accession number MT126808.1) was used in this work. The virus was grown in Vero cells in the Laboratory of Molecular Virology, at Federal University of Rio de Janeiro, Brazil. Vero cells were maintained in DMEM supplemented with 5% fetal bovine serum (FBS; Gibco) at 37 °C and 5% CO_2_. All work involving infectious SARS-CoV2 was performed in a biosafety level (BSL)-3 containment laboratory.

### Infection assays

Semi-confluent (70%) cells were grown on sterile glass coverslips in 24-well tissue culture plates infected with MOI (multiplicity of infection) values of 0.01, 0.1, or 1 using SARS-CoV-2 in free-serum medium. Fresh medium containing 5% FBS was added after an absorption period of 1.5 h at 37 °C and 5% CO_2_. Cells were processed for electron microscopy 2 or 48 hours post-infection (hpi).

### High Resolution Scanning Electron Microscopy

After 2 or 48 hours post-infection (hpi), samples were fixed with 2.5% glutaraldehyde in 0.1 M cacodylate buffer (pH 7.2) for 2 h. The coverslips were washed with 0.1 M sodium cacodylate buffer and post-fixed for 40 min in 1% OsO_4_ with 0.8% potassium ferrocyanide. After another washing cycle, the samples were dehydrated through a series of increasing concentration (30%–100%) of ethanol. The samples were critical-point-dried in liquid CO_2_ in a Balzers CPD apparatus before monolayer scraping with Scotch tape™ as previously described [18]. They were then sputtered with a 5-nm thick platinum coat in a Balzers apparatus. Samples were observed using an Auriga ZEISS microscope operated between 1.0 and 1.8 Kv with resolution of 3072-2304 and aperture size of 20 μm.

## 3. Results

To identify alterations on the surface of SARS-CoV-2-infected cells, we compared their morphology and the occurrence of surface projections (SP). While we did not detect any significant alteration in cell shape, the presence of SP increased on the surface of infected cells at 2 hpi (Fig. 1A-C). However, no viral particles were adhered to the cell surface or beneath these projections (Fig. 1D). At 48 hpi, we compared the surfaces of mock and infected cells (MOI of 0.1) to highlight the presence of viral particles adhered to the smooth cell surface and to the SP (Fig. 1E-F).

**Figure 1.**
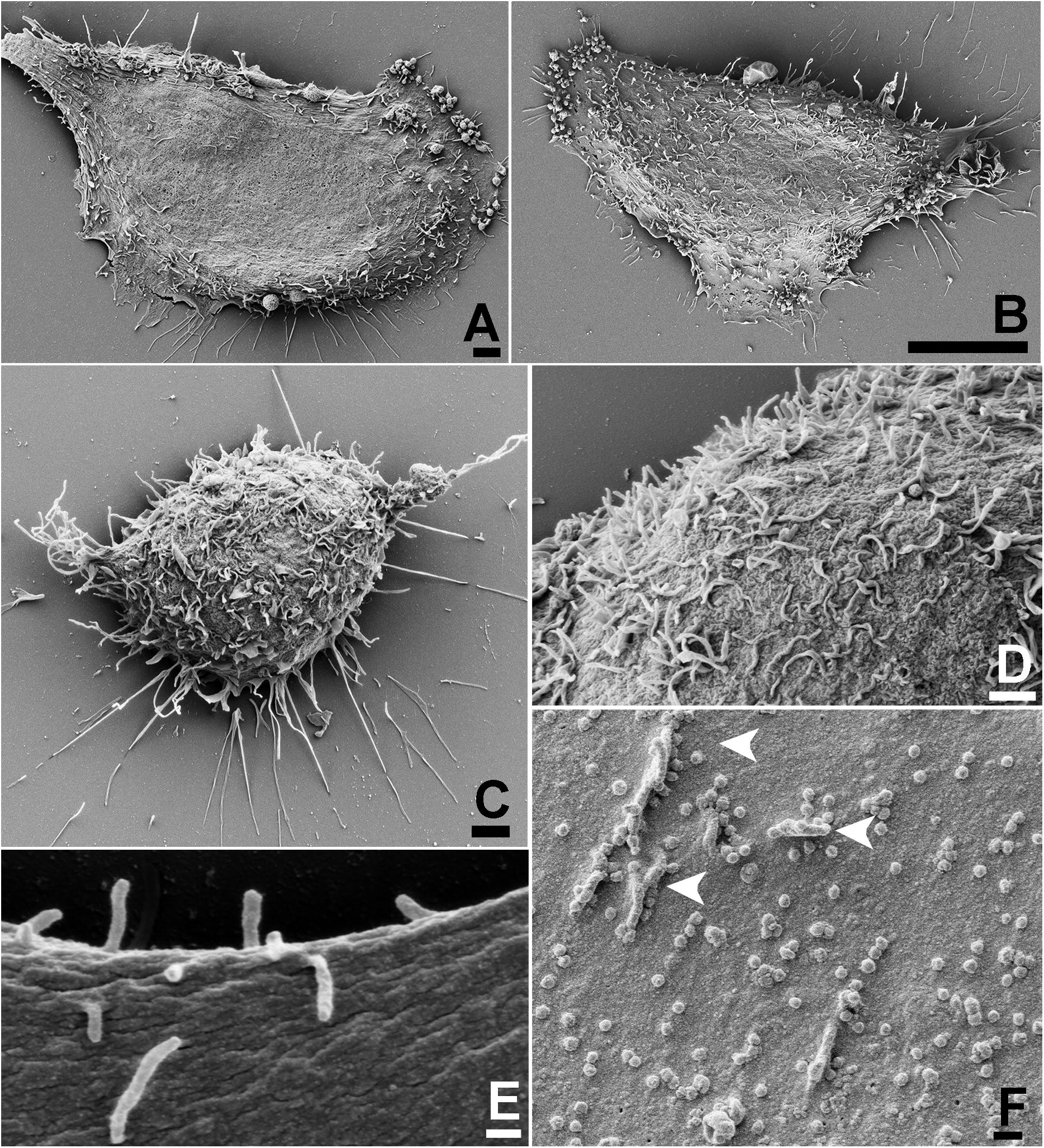
Effect of SARS-CoV-2 infection on host cell surface at 2 and 48 hpi. At 2hpi, mock-infected cells exhibited smooth surfaces **(A)**, while infected cells presented a discreet increment in the number of SP with the MOIs of 0.01 **(B)** and 1**(C)**. No viral particles were observed on the surface of infected cells at 2hpi, even with the MOI of 1 **(D)**. **(E)** Mock-infected cell surface at 48 h. **(F)** Virus adhesion to the cell surface and SP (arrowheads) became more evident with the MOI of 0.1 **(F)**. Bars: **(A,C)** 2 μm; **(B)** 10 μm; **(D)** 1μm; **(E-F)** 200 nm.

At the same time, and with a MOI of 1, viruses that egressed from a previous cycle of infection were observed during the process of attachment to the cell plasma membrane (Fig. 2A). The corona-like features of the SARS-CoV-2 particles were discernible via SEM (Fig. 2B), and the measurements showed sizes of approximately 80 nm in diameter (Fig. 2C).

**Figure 2.**
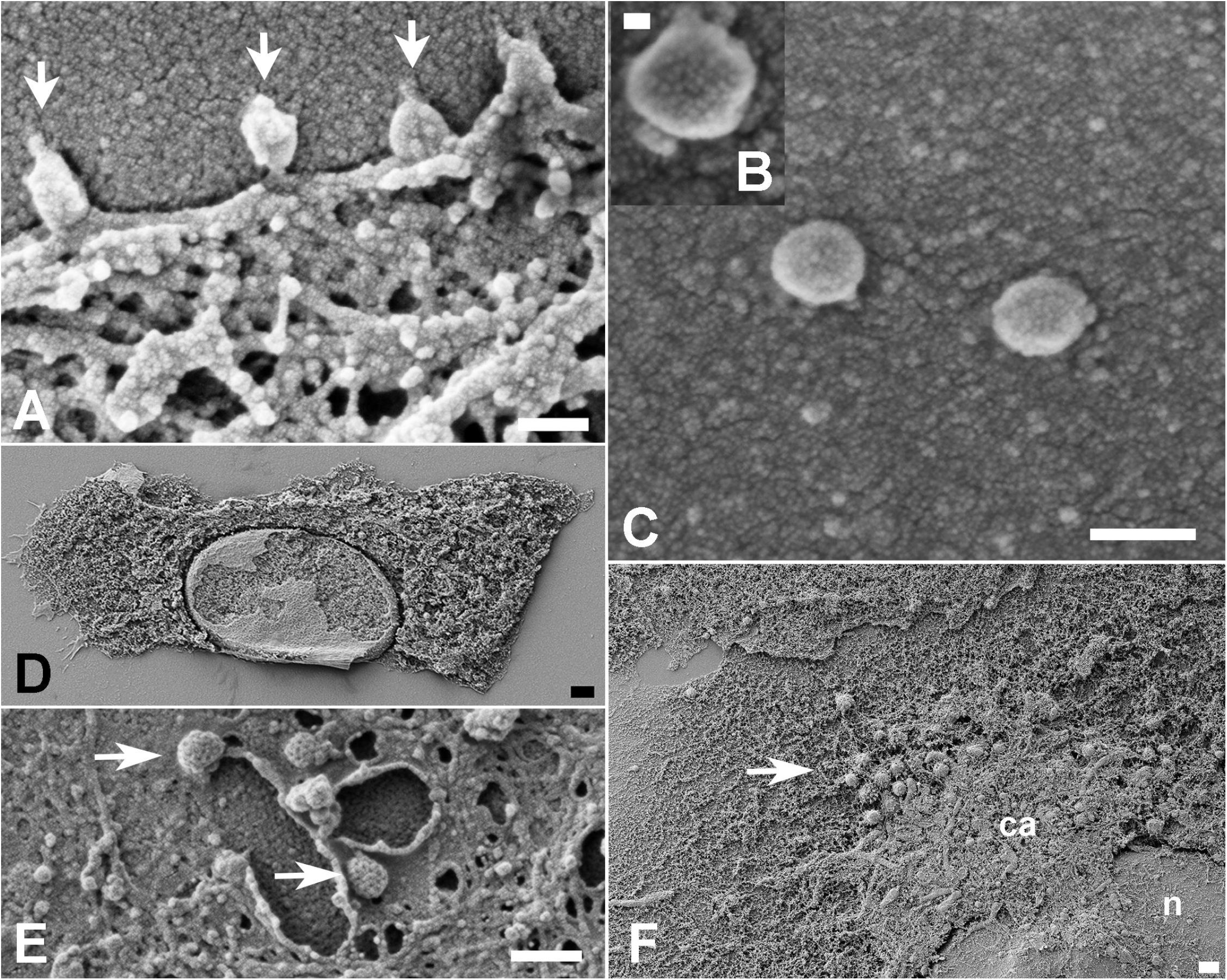
Morphology of cells surface and interior at 48 hpi. **(A)** With the MOI of 1, virus attachment (arrowheads) was frequent. **(B)** Spikes of SARS-CoV-2 particles (arrows) observed on the cell surface (cs) were discernible (MOI of 0.1). Viruses were observed laid on cell surface and adhered (arrow) to SP (mv) of infected cells. They exhibited a size between 70 and 85 nm **(B-C)** at MOIs of 0.01 and 0.1 respectively. Scraping of cells plasma membrane **(D)** revealed a homogeneous distribution of organelles in mock-infected samples, Infected cells exhibited coated pits vesicles of □ 100 nm (arrows) at perinuclear sites **(E)**. A polarized disposal at the infected cells cytosol **(E)** represented as a condensed area (ca) in the infected ones (MOI of 0.1). (n): nucleus; Bars: **(A, C, E)** 100 nm; **(B)** 20 nm; **(D)** 2 μm.

Removal of the host cell plasma membrane before platinum sputtering exposed the interior of the mock and infected cells. While mock cells displayed a diffuse distribution of organelles (Fig. 2D), infected cells exhibited a more polarized disposition of organelles and pit-coated vesicles approximately 100 nm in diameter (Fig. 2E-F). With a MOI of 1, cells at 48 hpi showed a plethora of vacuoles (0.4 to 1 μm; Fig. 3A). These were translocated to the cell plasma membrane presumably to perform exocytosis of viral particles (Fig. 3B). Some of these vacuoles had their content revealed and were filled with immature viruses, amorphous materials, or a hemocyte-like format (Fig. 3C-E). Although no virus-like particles could be distinguished in the ER, bordering vesicles were observed on the vacuoles (Fig. 3D).

**Figure 3.**
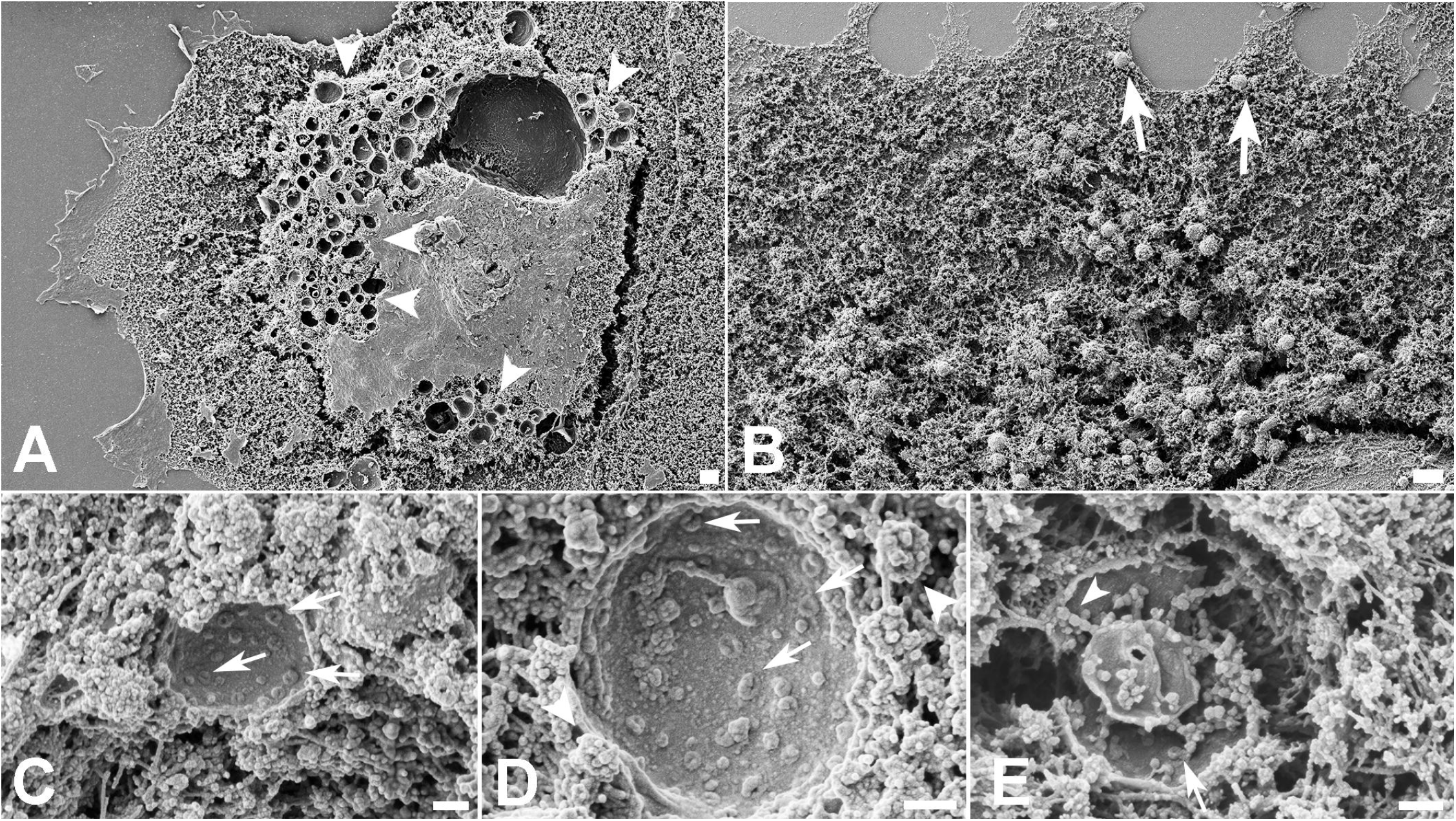
Inspection of the condensed areas of cells at 48 hpi. **(A)** Profusion of vacuoles (arrowheads) in cells infected with the MOI of 1. The possible route of the vacuoles was indicated by arrows in **(B)**. Scraped vacuoles had at least part of their content exposed. Vacuoles in **(C)** and **(D)** presented doughnut-like particles (arrows). In **(D),** borderinng vesicles (arrowheads) could be recognized next the vacuole membrane. Vacuoles shown in **(E)** display doughnut-like particles (arrow) and immature viral-like particles (arrowhead) too. MOIs: **(B-D)** 0.1; **(E)** 0.01; Bars: **(A-B)** 1 μm; **(C-D)** 200 nm.

Cells at 48 hpi also had viral particles near the cell surface membrane ruffles (Fig. 4A) and a filopodium-like structure (Fig. 4B). Other viral particles were wrapped with thin (□ 70 nm) cellular projections that resemble nanotubes (Fig. 4C). Membrane bridges that connect two cells showed the presence of virus particles on their surface (Figs. 4D-E).

**Figure 4.**
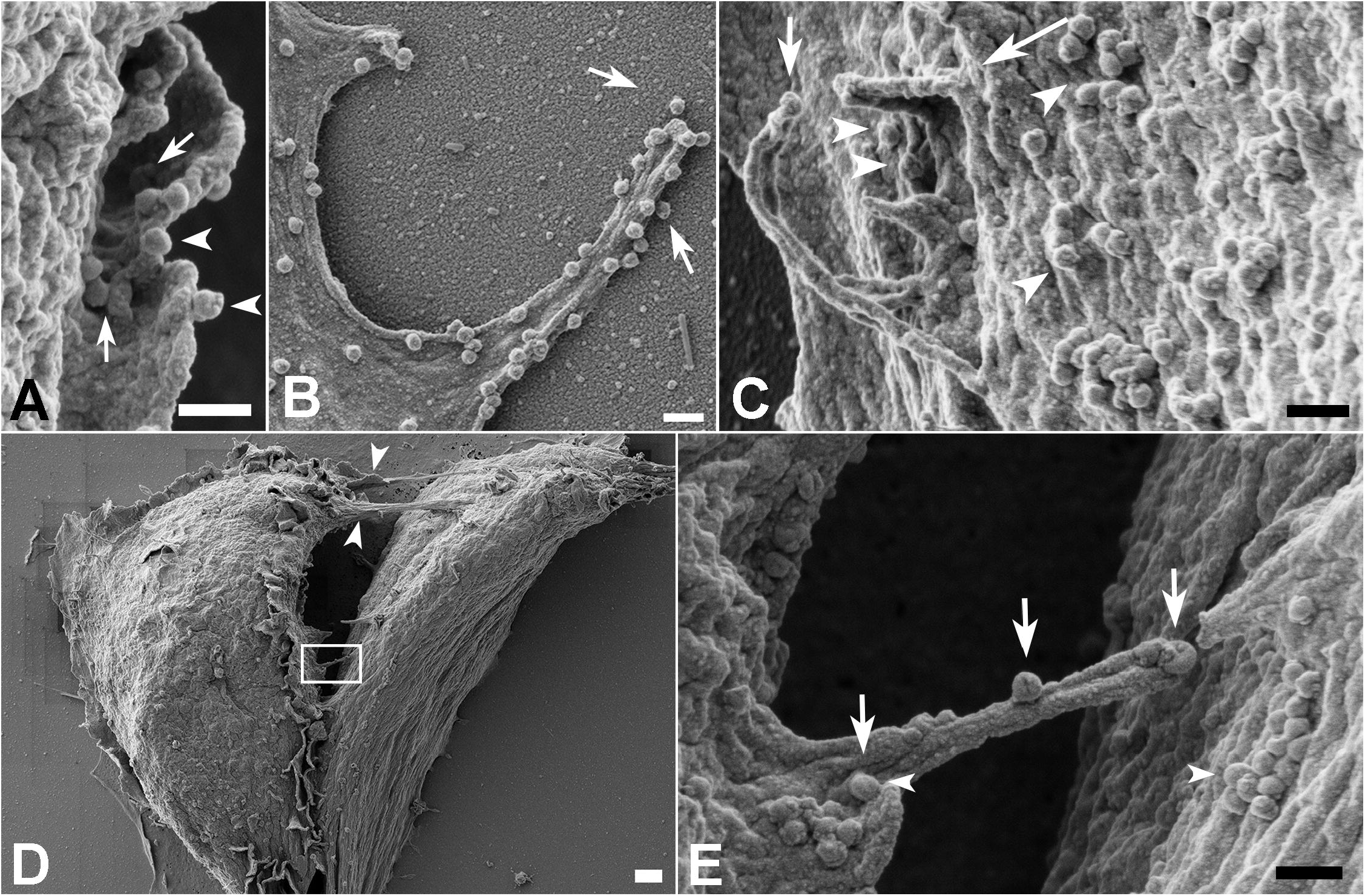
Fate of the SARS-CoV-2 particles adhered to the cell surface at 48 hpi. **(A-B)** Cell membrane ruffles about to wrap several viral particles (arrows). Viruses could also be observed on the edge of membrane ruffles (arrowheads) **(A)** and phyllopodium-like extensions **(B)**. In **(C)** a viral particle could be seen adhered to the edge of the microvilli-like structure (arrow) next to membrane ruffles (long arrow) surrounding SARS-CoV-2 particles (arrowheads). **(D)** Communications between two infected cells are indicated with arrowheads. One of the bridges between the cells was depicted in the rectangle. A higher magnification of this area is shown in **(E)** and displays viral particles (arrows) on their surface. Aggregates of SARS-CoV-2 particles (arrowheads) were also observed on the surface of both cells. MOI = 0.1; Bars: **(A-C, E)** 200 nm; **(D)** 1 μm.

## 4. Discussion

Part of the challenge in controlling COVID-19 is the innovative features of this coronavirus. New knowledge on virus genetics and morphology needs to be analyzed concurrently with viral “behavior” within the host cell as well as the dynamics that determine the fate of the particle. To approach SARS-CoV-2/cell interactions, we investigate several steps of virus infection at 2 and 48 hpi by SEM. This microscopic approach detailed virus-induced changes in the cell.

Our assays were performed using three MOIs (0.01; 0.1 and 1), and we could discern the MOI of 0.1 as the more adequate for this type of study. This MOI allowed the best cell conditions and distribution and also allowed visualization of virions through the cell surface into the cell interior.

The absence of virions adhered to the cells surface at 2 hpi corroborates recent studies performed by Belhaouari et al. [19] in which SARS-CoV-2 particles were only observed at these loci after 12 hpi. In contrast, SARS-CoV-2 particles were found lying on the cellular surface at 48 hpi between surface projections and adhered to them. We also observed probable viral particles inside vacuoles suggesting a secretion route. These aggregates of cell organelles and components (Fig. 2F) may reflect the polarized release of virus previously described for SARS-CoV [20].

All viruses measured by SEM display a spiky round shape with a size of around 70-85 nm in diameter considering a platinum coating of 5 nm. This agrees with the dimensions described in recent studies [1,21,22].

### 4.1 Viral particles adhered to smooth surface and microvilli-like surface projetions

The effects on the surface morphology of infected cells varies among viruses. Infection by several viruses including HTLV-III_B_ leads to a loss of cell SP that are then replaced by blebs [23]. Microvilli induction or increases were reported in several cases of DNA or RNA viral infection [24,25]. For RNA viruses that egress by budding, e.g., influenza, the increase in SP of infected cells coincides with higher budding rates [26].

Similar to prior studies on SARS-CoV infection of Vero cells [27], we also observed a ruffled host cell and thickened edges displaying a layered shape. These sites were appropriate to register the attachment of SARS-CoV-2 particles (Fig. 2A) similar to transmission electron microscopy images of the same early step of SARS-CoV infection of Vero cells [28].

Likewise, the proliferation of SP on the infected cells, especially at the apical region of these cells, is similar to SARS-CoV and SARS-CoV-2. In addition, the abundance of SARS-CoV-2 particles held on SP may facilitate the speed of viral propagation in the epithelium of conducting airways from the lumen of the respiratory superior tract because this environment is colonized by ciliated cells.

### 4.2 Vacuoles containing viral particles

Cell scraping is a very useful expedient that is occasionally used in studies of host cell/parasite interactions [29,30]. Infected cells are artificially devoid of plasma membranes and exposed to a myriad of vacuoles (Fig. 3A). Drastic vacuolization due to viral infection was previously described for other RNA viruses including SARS-CoV [20,31]. Similar sites were recently reported as virus morphogenesis matrix vesicae (VMMV) [19]. The particles observed in the interior of these VMMVs (Fig. 3C-E) were previously described as doughnut-like particles when observed by electron microscopy [32,19]. SARS-CoV immature particles are presumed to bud into vesicles as part of the assembly process, and thus the observed particles were probably immature viruses devoid of the representative (corona) spikes of this virion. Bordering vesicles were found in close association with the vacuoles (Fig. 3D), and thus we speculate that their role in viral pre-components leads to discharge into the compartments.

Studies with other coronaviruses identified large virion-containing vacuoles (LVCVs) where the complete particle would bud. There is correlation between these structures as observed by transmission electron microscopy and our data suggesting the occurrence of both phenomena.

### 4.3 Translocation of vacuoles towards the plasma membrane

Coronaviruses infection leads to massive remodeling of cell membranes [33,34]; the more condensed area depicted in the cytoplasm at 48 hpi (Fig. 2F) may correspond to the main locus of viral morphogenesis. The proposed mechanism for the export of viruses to the extracellular space is via fusion of the transport compartment membrane with the cell plasma membrane [20].

The size of the vacuoles we observed in the cell periphery was not compatible with the identified clathrin-coated pits because the vacuoles measure approximately 1 μm; clathrin-coated pits measure near 200 nm in diameter. The presence of these endocytosis-associated players was recently reported in SARS-CoV-2-infected cells. They are likely receptacles to the nucleocapsid after the incoming virus is uncoated [19].

Thus, our observations suggest that a boost in vacuoles is restricted only to a specific and more condensed part of the cytoplasm. This suggests translocation to the plasma membrane is required for release the viral particles by a fusion mechanism.

### 4.4 Cellular bridges containing viral particles

Viral particles adhered to cell surface protrusions that were shown to connect two cells. This observation suggests viral “cell surfing” previously described by other enveloped viruses such as HIV and human metapneumovirus [35,36]. This mechanism is presumed to allow the *in vivo* penetration of virus in mucosal surfaces that display microvilli-rich cells.

Actin filaments play a fundamental role in viral extrusion by the cell for both RNA and DNA viruses. Actin offers the strength to discharge the progeny virus particles to the extracellular medium, as occurs to some viruses that leave the cell by budding, including Fowlpox and West Nile viruses [37,38]. Other examples include actin comets—these are an efficient form of poxvirus dissemination and cell-to-cell HIV spreading, which involves the direct engagement of GAG proteins and F-actin [39,40].

Previous studies have shown that the cytoskeleton network plays an important role in the maturation and, possibly, in the replication process of SARS-CoV [27]. Communication between the two cells in Fig. 4C-D suggests the occurrence of a thin (< 0.7 μm) strand of Factin containing tunneling nanotube (TNT). These intercellular membranous connections may provide the transference of molecular information especially viruses [41]. Similarly, virus cell surfing was shown on SARS-CoV-2 infection, which offers new insights into cell-to-cell propagation and virus transmission.

## Data availability

The data that support the findings of this study are available from the corresponding author on reasonable request.

## Acknowledgements

We thank Dr. Lorian Cobra Straker for technical assistance. This work has been supported by Fundação Carlos Chagas Filho de Amparo à Pesquisa do Estado do Rio de Janeiro-FAPERJ, Financiadora de Estudos e Projetos-FINEP and Conselho Nacional de Desenvolvimento Científico e Tecnológico-CNPq.

